# Dynamics of infection in a novel group of promiscuous phages and hosts of multiple bacterial genera retrieved from river communities

**DOI:** 10.1101/2020.08.07.242396

**Authors:** Daniel Cazares, Adrian Cazares, Wendy Figueroa, Gabriel Guarneros, Robert A. Edwards, Pablo Vinuesa

## Abstract

Phages are generally described as species- or even strain-specific viruses, implying an inherent limitation for some to be maintained and spread in diverse bacterial communities. Moreover, phage isolation and host range determination rarely consider the phage ecological context, likely biasing our notion on phage specificity. Here we identified and characterized a novel group of promiscuous phages existing in rivers by using diverse bacteria isolated from the same samples, and then used this biological system to investigate infection dynamics in distantly related hosts. We assembled a diverse collection of over 600 native bacterial strains and used them to isolate six podophages, named Atoyac, from different geographic origin and capable of infecting six genera in the Gammaproteobacteria. Atoyac phage genomes are highly similar to each other but not to those currently available in the genome and metagenome public databases. Detailed comparison of the phage’s infectivity in diverse hosts and trough hundreds of interactions revealed variation in plating efficiency amongst bacterial genera, implying a cost associated with infection of distant hosts, and between phages, despite their sequence similarity. We show, through experimental evolution in single or alternate hosts of different genera, that plaque production efficiency is highly dynamic and tends towards optimization in hosts rendering low plaque formation. Complex adaptation outcomes observed in the evolution experiments differed between highly similar phages and suggest that propagation in multiple hosts may be key to maintain promiscuity in some viruses. Our study expands our knowledge of the virosphere and uncovers bacteria-phage interactions overlooked in natural systems.

**Importance:** In natural environments, phages co-exist and interact with a broad variety of bacteria, posing a conundrum for narrow-host-range phages maintenance in diverse communities. This context is rarely considered in the study of host-phage interactions, typically focused on narrow-host-range viruses and their infectivity in target bacteria isolated from sources distinct to where the phages were retrieved from. By studying phage-host interactions in bacteria and viruses isolated from river microbial communities, we show that novel phages with promiscuous host range encompassing multiple bacterial genera can be found in the environment. Assessment of hundreds of interactions in diverse hosts revealed that similar phages exhibit different infection efficiency and adaptation patterns. Understanding host range is fundamental in our knowledge of bacteria-phage interactions and their impact in microbial communities. The dynamic nature of phage promiscuity revealed in our study has implications in different aspects of phage research such as horizontal gene transfer or phage therapy.

## Introduction

Bacteriophages, or phages for short, have been systematically described as species- or even strain-specific viruses (1–3). Since phage infection is initially conditioned to chance encounters with a receptive host cell (4), the premise of the phages narrow-host-range implies a severe limitation for them to be maintained and spread within environments featuring diverse microbial communities (4), which could promote the existence of phages with a wider range of potential hosts. Although phages capable of infecting bacteria beyond the taxonomic rank of species or genus have been previously described, these are considered rare when compared to the far larger number of phages reported as specific to a single bacterial species (1–3).

Characterization of the host range is a paramount step towards assessing phage specificity and identifying specimens with a broad scope of host targets. Since the host range of a bacteriophage is described as the taxonomic diversity of hosts it can successfully infect (5), the strategies used to interpret a “successful infection” are key to characterize this trait. In order to infect, phages must attach to the host cell, transport their genomes into the host cell, overcome various defense mechanisms to hijack the host’s molecular machinery in order to replicate, assemble and release the new viral progeny (5, 6). Infection assays in liquid cultures, known as lysis curves, and plaque assays resulting in the formation of either lysis spots or individual lytic plaques, represent examples of commonly-used methods to infer phage infection (2). As factors other than phages (e.g. pyocins) and mechanisms independent of phage production (e.g. lysis-from-without) can also lead to cell death by lysis, the isolation of individual phage lytic plaques is regarded as the best approximation to identify productive infection, i.e. an infection cycle yielding new phage progeny (2).

In spite of being labeled as infrequent, the isolation and study of phages with the ability to infect multiple species or genera, can significantly contribute to advancing our understanding of the host range evolution and phage ecology. Recent metagenomics studies have provided valuable insights into the distribution of broad-host-range phages in different environments (5, 7, 8), however, the nature of this approach restricts further experimental characterization of these viruses which could target unculturable bacteria or strains that have not been isolated yet. On the other hand, some pioneering studies have focused on the targeted isolation of broad-host-range phages through enrichment procedures by either co-culture (9) or consecutive monoculture (10) with potential hosts of different taxonomic origin. Such approaches led to the isolation of interesting unrelated specimens infecting bacteria from multiple species including *P. aeruginosa* and *Escherichia coli*.

Here we addressed the identification of broad-host-range phages by assembling a large collection of bacterial and phage isolates present in freshwater and wastewater samples. We hypothesized that using a taxonomically-diverse panel of bacteria native to these environments as potential hosts would facilitate the isolation of phages infecting multiple species from such microbial communities. This approach led to the identification of a group of closely-related podophages, named Atoyac (Nahuatl for river), capable of replicating in bacteria from six different genera and three orders within the Gammaproteobacteria class. These phages, here referred to as “promiscuous”, comprise a novel viral group with undetectable presence in available metagenomic data. The Atoyac phages display different plating efficiency amongst the genera they infect, however, such efficiency rapidly changes over consecutive propagation in a single or multiple genomic backgrounds, indicative of dynamic host adaptation. The adaptation trajectories are phage-dependent and lead to different outcomes despite the high similarity existing between the phages. Our findings contribute to expanding the virosphere and provide insights into short-term host-range evolution and virus-host interactions in nature.

## Results

### Isolation of promiscuous phages

We searched for promiscuous phages, i.e. phages capable of infecting hosts of distinct taxonomic origin, by assembling a large and diverse collection of over 600 bacterial isolates and free phages existing in wastewater and environmental water samples from twelve different sites in Mexico (Supplementary Figure 1, see Methods). Infectivity assays showed most of the water samples’ phages produced plaques when spotted on bacterial lawns of the panel of native isolates generated from the sampled sites. Importantly, evaluation of phage-bacteria interactions through plaque assays in standard media such as TΦ or LB did not always allow identification of phage infectivity, however, exploring other conditions and diverse media (see Methods) led to formation and identification of phage plaques. A cross-infection screening amongst the susceptible isolates led to the identification of phages producing lytic plaques in several strains with distinct phenotypes (Supplementary Figure 1). Remarkably, six phages isolated from five sites and named Atoyac were able to infect 54 isolates in the panel. Susceptible bacteria included members of the genera *Aeromonas, Pseudomonas, Hafnia, Escherichia, Serratia*, and *Yersinia*, confirming the promiscuous nature of the Atoyac phages’ host range, which crosses multiple species barriers within the Gammaproteobacteria class (Figure 1, Supplementary Figure 2). The Atoyac phages were isolated from river and sewage water samples obtained from Central, Southeast, and Northwest regions in Mexico (Supplementary Figure 3). Assessment of fecal contamination by either coliform count or presence of crAssphage (used as a marker to detect human fecal pollution in water (11)), revealed that the Atoyac phages were prevalent in contaminated sites which represented most of our samples (Supplementary Table 1). Contaminated samples were also the source of the highest phage titers determined in the infection assays.

**Fig. 1.**
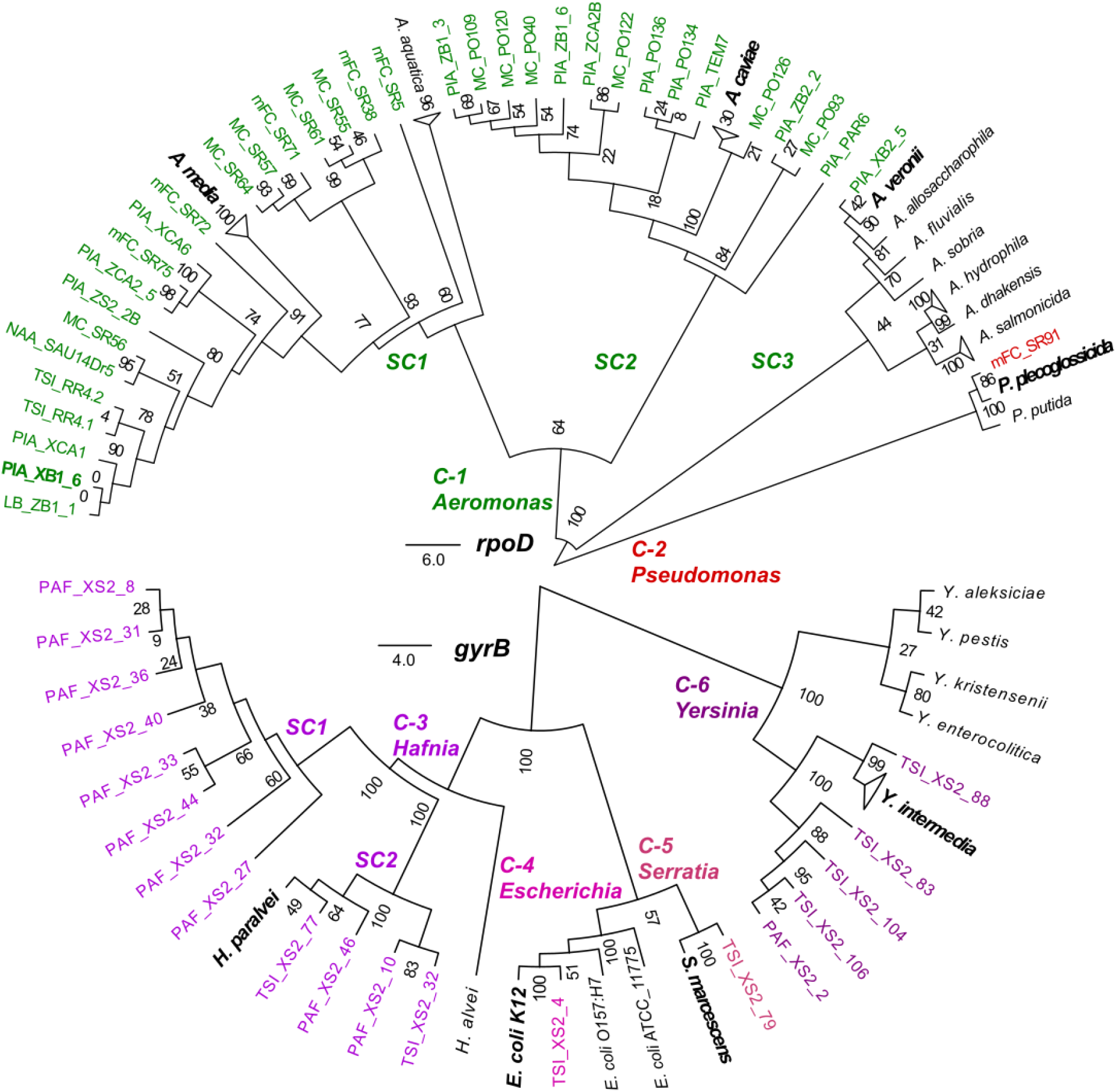
Environmental isolates susceptible to infection by the promiscuous Atoyac phages. Neighbor-joining trees of the bacterial isolates infected by phages of the Atoyac group were reconstructed from the alignment of nucleotide sequences of the genes *rpoD* (top) or *gyrB* (bottom) under the J-C substitution model. Strains were considered as susceptible only when Atoyac phages formed individual lytic plaques on them (See Methods). The 54 isolates from our collection are indicated in colour labels corresponding to the bacterial genus they belong to. Reference strains of different bacterial species included in the trees are indicated in black labels, and those clustering with isolates from our collection are highlighted in bold letters. Clusters and sub-clusters of the different bacterial genera identified in the tree are indicated with the “C-” and “SC” prefixes, respectively. Branch support values from 100 replicas bootstrap tests are shown in the trees. The scale-bar represents the expected number of substitutions per site under this model.

### Efficiency of plating of the Atoyac phages

The six Atoyac phages displayed a similar host range, with plaque morphologies varying depending on the host group and being generally larger in *Aeromonas* isolates (Supplementary Figure 2). Notable between-phage variation in plaque morphology was observed in lawns of the *Pseudomonas* isolate. Differences in host range detected amongst the phages corresponded to the ability to infect some of the *Aeromonas, Hafnia*, and *Yersinia* isolates (Figure 2A). Phage Atoyac15, for instance, was the only member of the group capable of infecting all the *Aeromonas* isolates in the panel, but could not infect most of the *Yersinia* strains, nor some representatives of the *Hafnia* group.

**Fig. 2.**
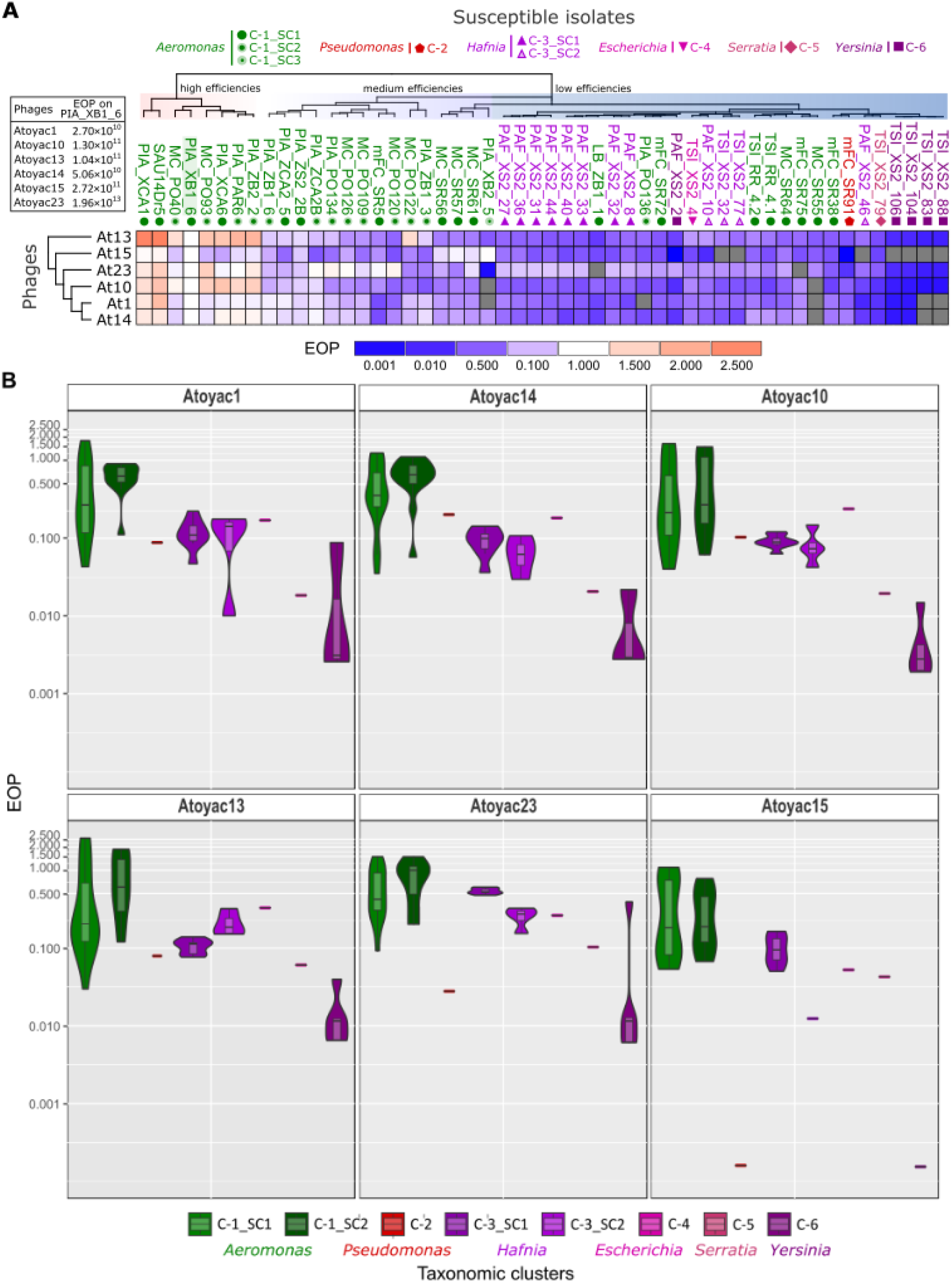
Host range and efficiency of plating of the Atoyac phages. **A.** The heatmap illustrates the Efficiency of Plating (EOP) recorded for six Atoyac phages (rows) on a panel of 54 environmental strains isolated in this work (columns). Names of the bacterial isolates are colour coded according to the taxonomic genus they belong to as indicated at the top of the figure. Symbols below the isolate names indicate the bacterial cluster or sub-cluster inferred through sequence alignment of a marker gene (see Fig. 1). The titer determined for each phage on the propagating strain PIA_XB1_6, indicated in the left inset, was used as the reference value to calculate the EOP. The scale at the bottom of the heatmap depicts the deviation from the plating efficiency recorded in the reference isolate (EOP = 1.0). Cells in grey correspond to interactions that did not generate detectable lytic plaques thus indicating non-susceptible strains. Both bacterial isolates and phages are hierarchically clustered based on the EOP values using the Euclidean distance method. **B.** Plotted are the EOP values recorded for each Atoyac phage grouped by the taxonomic cluster (genus) or sub-cluster of their hosts. Clusters are colour coded as in the figure panel A and Fig. 1. The EOP values were calculated from five independent biological replicates and the averages are plotted. Values in the violin plots are in log scale.

We determined the efficiency of plating (EOP) (12) across the panel of susceptible strains to gain further insights into the infection capacity of the Atoyac phages, and compared the EOP on each strain to the number of plaques formed on the reference strain *Aeromonas* sp. PIA_XB1_6 (Figure 2). The results revealed complex EOP profiles among the susceptible isolates, although a clear pattern emerged. We found that clustering the bacterial isolates based on the phages’ EOP largely reflects their taxonomic relatedness at the genus level (Figure 2A), suggesting that the efficiency of the Atoyac phages to produce Plaque-Forming Units (PFU) is influenced by the taxonomic origin of the host and that crossing the genus barrier imposes a cost. In general, Atoyac phages generated significantly more lytic plaques on *Aeromonas* isolates, followed by strains from the *Hafnia* and *Yersinia* genera, respectively (Figure 2, Supplementary Table 2). No statistically significant differences in plating efficiency per phage were detected between *Aeromonas* subclusters (Mann-Whitney U test and *p*-value 0.05), indicating that the major EOP differences occur between genera (Figure 2B, Supplementary Table 2). Some *Aeromonas* isolates featuring EOP values lower than those recorded in strains of other genera were also detected.

The EOP values measured in the collection of susceptible strains ranged from −1000 to 2.5 times the EOP on the reference strain *Aeromonas sp*. PIA_XB1_6 (original propagative strain, see Methods). For example, the stock of the phage Atoyac13 produced 6.72 x 10^8^ PFU/mL on a bacterial lawn of *Yersinia sp*. TSI_XS2_88, 2.68 x 10^11^ PFU/mL on *Aeromonas sp*. PIA_XCA1, and 1.04 x 10^11^ PFU/mL on the reference strain *Aeromonas sp*. PIA_XB1_6 (Supplementary Table 2).

### Virion morphology and genome comparison

We characterized the Atoyac phages at the virion morphology and genome levels to investigate the similarities between them and to other phages. Electron microscopy analysis of the CsCl-gradient purified viral particles revealed a shared morphology typical of phages of the *Podoviridae* family, featuring a ~50 nm icosahedral head attached to a very short tail (Figure 3, Supplementary Figure 4).

**Fig. 3.**
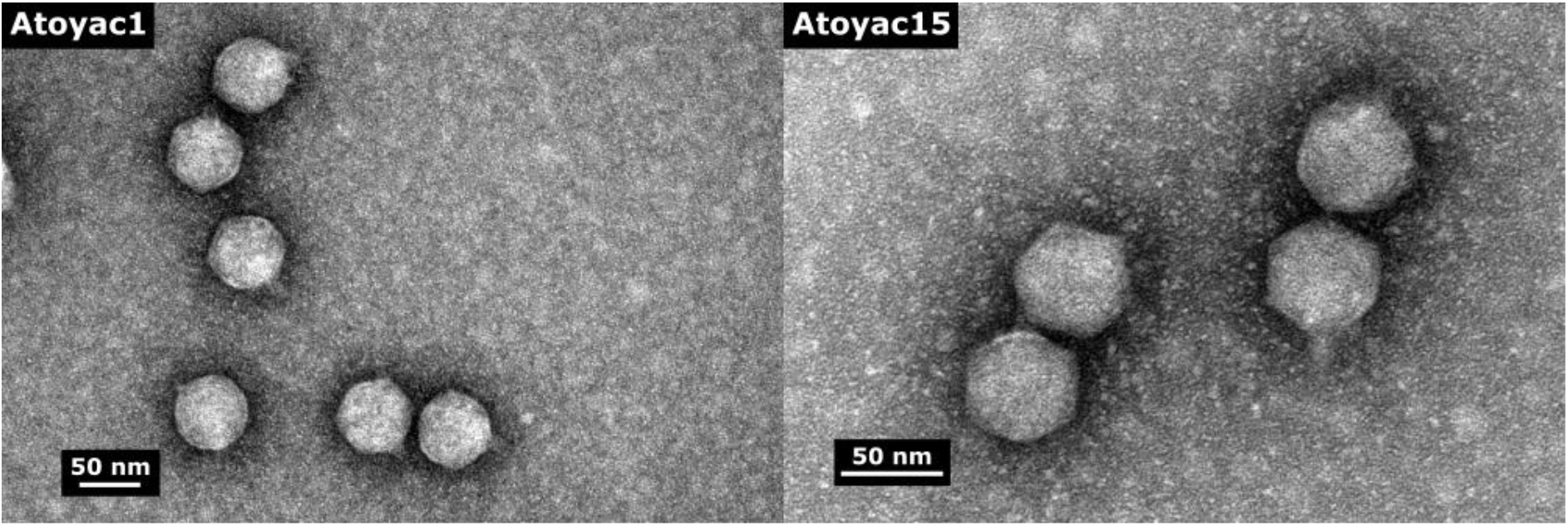
Electron micrographs of virions of the Atoyac phages 1 and 15. The CsCl-purified viral particles were negatively stained with 2% uranyl acetate and visualized at 150000-fold magnification. Rod on the micrographs represents 50 nm size.

The Atoyac phage genomes average 41.7 kb in size and 59% GC content. A comparative analysis unveiled that they comprise a monophyletic group sharing from 85.9 to 98.4% overall nucleotide sequence identity (Figure 4, Supplementary Figure 5). Interestingly, we found a good correlation between the clustering of the Atoyac phages based on the whole genome comparison (Supplementary Figure 5) and that inferred from the EOP and host range profiles (Figure 2A), suggesting that sequence variability largely accounts for the differential infection capability displayed by the phages. We identified two sub-groups of phages (Group 1: phages 1,14 and 10, Group 2: phages 13 and 23) that were more closely related to each other in the whole genome tree and displayed very similar infection profiles. Phage Atoyac 15 was the most divergent in the genome comparison and also exhibited one of the most dissimilar infection profiles. The major source of diversity among the Atoyac phage genomes corresponded to a large noncoding region featuring GC content lower than the global average and an adjacent cluster of ORFs encoding mostly small (< 100 aa) hypothetical proteins of unknown function (Figure 4, Supplementary Figure 6). Marked sequence variation was also detected in contiguous ORFs putatively encoding a tail fiber and a glycosidase superfamily protein.

**Fig. 4.**
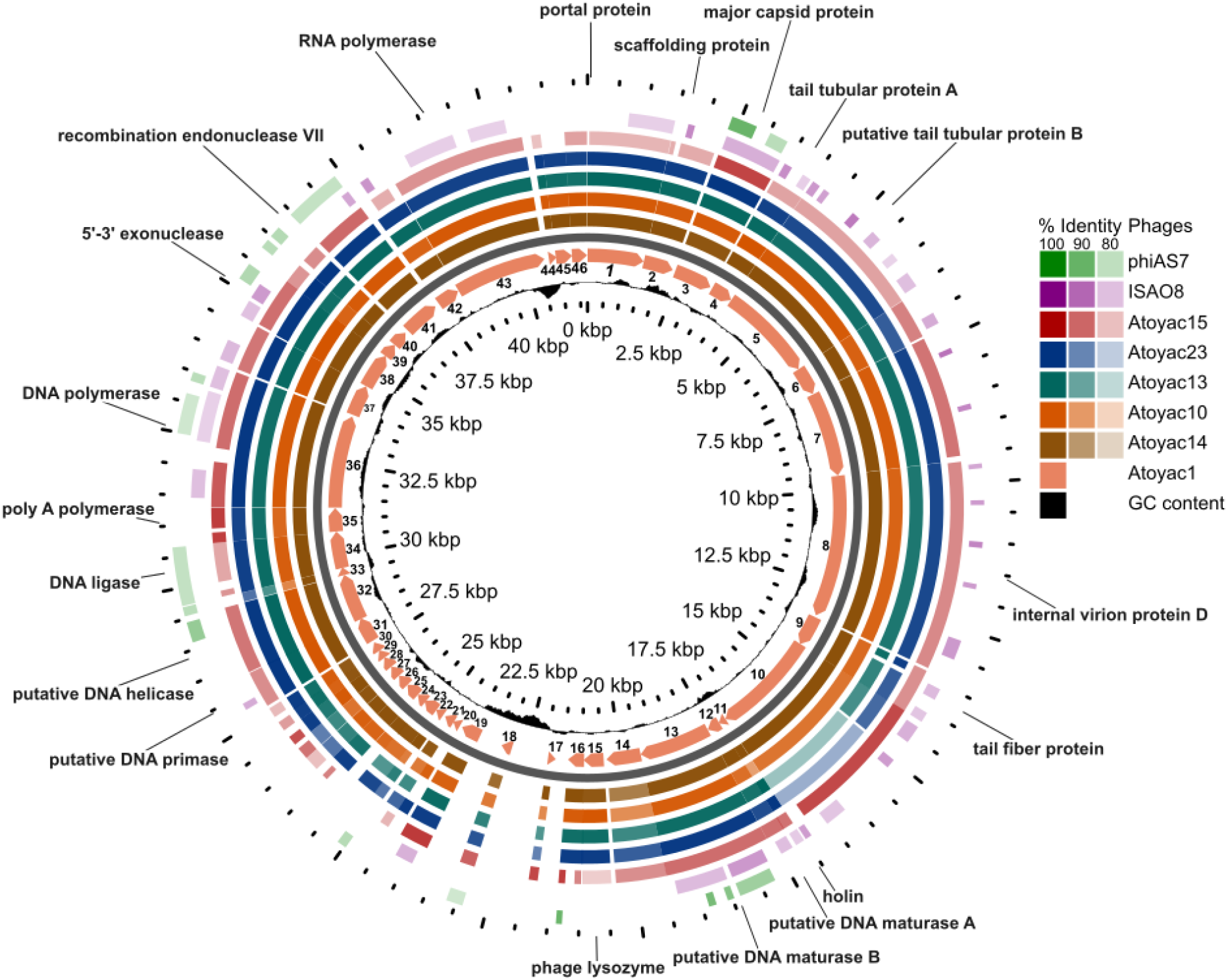
Comparative genomics analysis of the Atoyac phage group. Genomes of six Atoyac phages and two distant homologues are represented as colour rings and indicated at the right side of the figure. The genome of phage Atoyac 1, represented with a map in the innermost colour ring, was used as a reference to compare the ORF sequences of the other phages at nucleotide level. Arrows in the map correspond to ORFs pointing towards the direction of their transcription. Functions identified in the genome are indicated outside the rings pointing to the corresponding Atoyac 1 ORFs. Coordinates of the reference genome and the distribution of its GC content regarding the average (59%) are indicated inside the corresponding ring. The level of sequence identity detected in the genomes (>79%) respecting the Atoyac 1 ORFs is colour coded and indicated in the figure. The two outermost rings correspond to representatives of the phage groups displaying the highest sequence similarity to genomes of the Atoyac group.

Of the 46 to 50 ORFs identified in the Atoyac genomes, 23 were assigned with a function (46 to 50%) based on sequence homology and the presence of conserved domains. These ORFs are organized in two major functional modules encoding proteins related to: 1) virion morphogenesis and release such as capsid, tail and lysis proteins; and 2) transcription and replication, including RNA and DNA polymerases, a DNA primase, helicase, ligase, exonuclease and endonuclease with a potential role in recombination (Figure 4). Genes associated with a temperate lifestyle (e.g. repressor or integrase) were not identified in the genomes, implying that the Atoyac phages are virulent. This observation is consistent with our unsuccessful attempts to isolate lysogens of Atoyac phages in the *Aeromonas* and *Yersinia* backgrounds.

Homology searches revealed that the Atoyac phages constitute a novel group. Database searching revealed two groups of phages distantly related to Atoyac clade. The most similar representatives from these groups correspond to the podophages 4vB_YenP_ISAO8 infecting *Yersinia enterocolitica* (hereafter referred as to ISAO8, **NC_028850.1**) and phiAS7 infecting *Aeromonas salmonicida* (**JN651747.1,** (13)), which share 47.78 and 21.05% overall sequence identity with the genome of the phage Atoyac1, respectively (Figure 4, Supplementary Figure 5). We also compared the Atoyac phage genomes to that of the *E. coli* phage T7, a virulent archetype of the *Podoviridae* family. Despite the lack of extensive sequence similarity between the genomes at the nucleotide or protein levels, the presence of a large non-homologous and noncoding region was recognized in similar relative genome positions. A similar region has been identified in genomes of T7-like phages of different bacterial species and reported to contain multiple transcriptional promoters (14). In line with this observation, we identified a series of putative promoters in both strands of the large noncoding region of the Atoyac genomes (Supplementary Table 3, Figure 4).

### Search of Atoyac-like phages in metagenomics data

Atoyac phages were identified in different geographical regions in Mexico separated by ~3.9 to 2,525 km from each other, thus implying a broad distribution for this novel group of promiscuous viruses (Supplementary Figure 3). Since no other members of the group could be detected through homology searches in public genome databases, we decided to further investigate the distribution of the Atoyac phages by searching for similar sequences in available metagenomics data.

We sought to recover sequencing reads homologous to the Atoyac 1 genome by analysing metagenome datasets deposited in the NCBI sequence read archive (SRA). Surprisingly, our analysis of more than 65,000 datasets using a recently reported searching strategy (see Methods, (15)) did not identify metagenomes with sequencing reads matching the Atoyac reference. This result contrasts with those obtained when genomes of other two unrelated phages, crAssphage and T4, were used as reference and control for the search. Thousands of reads covering up to 100 and 86.5% of the crAssphage and T4 genomes were recruited in multiple samples (Supplementary Table 4), suggesting absence or low abundance of Atoyac-like phages in the analysed datasets. Alternatively, under-exploration of environments inhabited by Atoyac-like phages (e.g. fresh waters) through metagenomics approaches could account for this outcome (8).

### Experimental evolution of plating efficiency

Since the Atoyac phages displayed higher EOP in isolates of the *Aeromonas* group than in strains of the other susceptible genera, we took an experimental evolution approach to investigate the feasibility of reversing this trend in hosts with low plaque production efficiency. To diversify the set of hosts in the experiment, we chose *Yersinia* as representative of the *Enterobacteriaceae* family and *Pseudomonas* as the only *non-Enterobacteriaceae* genus identified as susceptible, besides *Aeromonas*. Isolates from the two selected genera recorded some of the lowest EOP values in the collection (Figure 2A). Phages 1 and 23, displaying distinct plating efficiency in the bacterial isolates chosen for the experiment (*Pseudomonas sp*. mFC_SR91 and *Yersinia sp*. PAF_XS2_2, Figure 2A), were selected as representatives of the two sub-clusters identified within the Atoyac group (Supplementary Figure 5).

Three individual plaques of Atoyac 1 and 23, each representing a lineage, were serially propagated for ten passages in a single host, *Pseudomonas* or *Yersinia* (see experimental design in Supplementary Figure 7, schemes A and B). To assess changes in plaque production over the course of the experiment, the EOP in *Pseudomonas* and *Yersinia* was calculated before the experiment and at the propagation steps 5 and 10, using *Aeromonas* as reference. As expected, the ancestral stock of both phages produced more plaques in the *Aeromonas* strain than in the isolates of the other two genera (Figure 5). Nevertheless, this difference in plating efficiency narrowed down in both *Pseudomonas* and *Yersinia* when the phages were consecutively replicated in the *Pseudomonas* isolate, suggestive of a cross-adaptation effect (Figure 5, panels A and B). A similar outcome was observed when Atoyac 1 was serially propagated in the *Yersinia* isolate (Figure 5, panel C), however, Atoyac 23 did not show a substantial change in EOP in the *Yersinia* strain and considerably reduced its virulence towards the *Pseudomonas* isolate (decrease in plaque number and size) under the same propagation conditions (Figure 5, panel D).

**Fig. 5.**
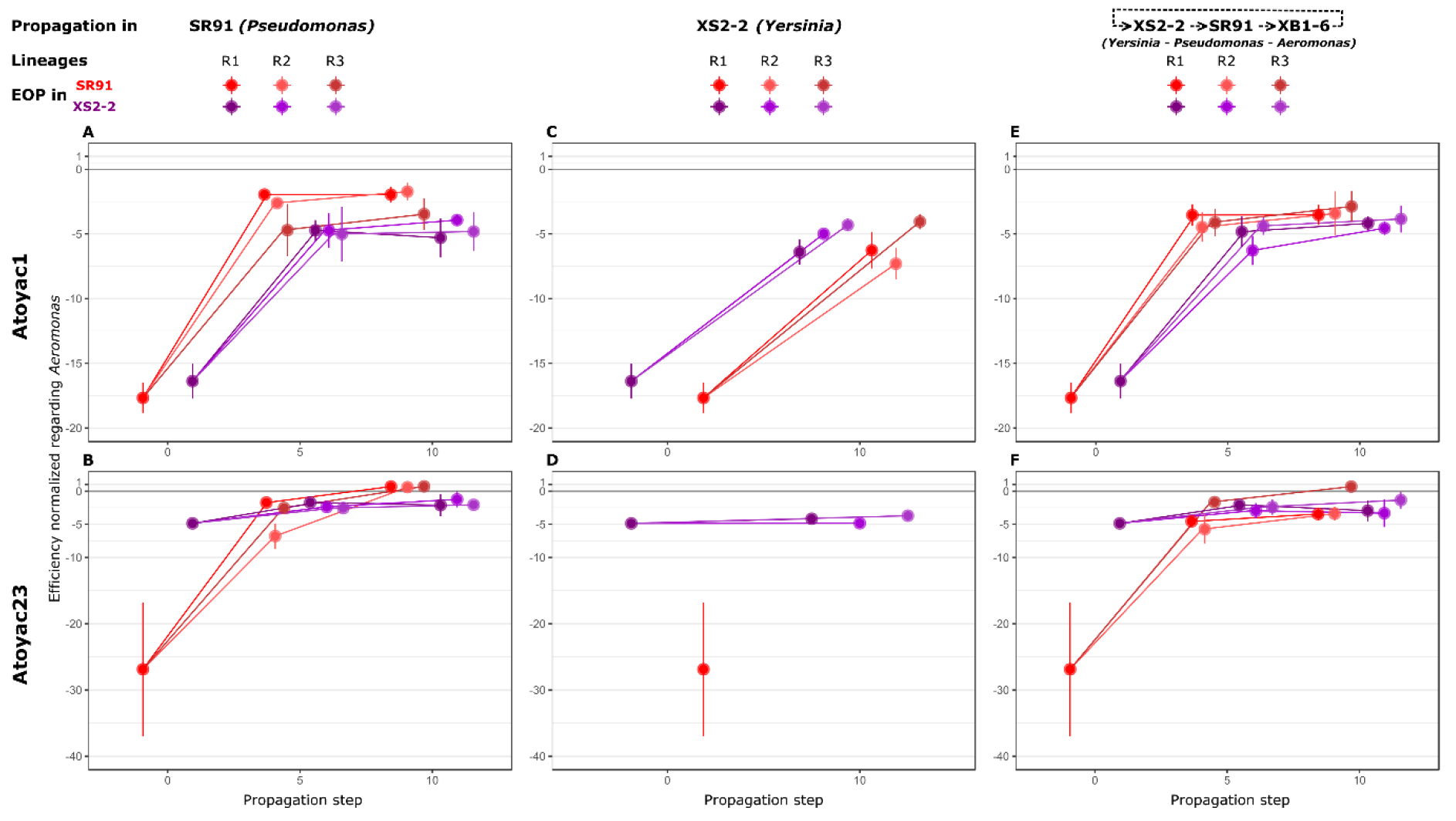
Experimental evolution of the efficiency of plating of two Atoyac phages. The charts represent the trajectories of the dynamics of efficiency of plating during experimental evolution of phages Atoyac 1 and 23. The upper row (sections A, C and E) shows the outcomes of the different propagation schemes used during the experimental evolution for the phage Atoyac 1. The lower row (sections B, D and F) illustrates the results for the phage Atoyac 23. In the first column (A and B), the charts show the difference in EOP between *Aeromonas* (XB1-6 as reference strain) and *Pseudomonas* SR91 (in red) or *Yersinia* XS2-2 (in purple) when the phages were serially propagated in *Pseudomonas* for 0, 5 and 10 passages. The values plotted in the propagation step 0 correspond to the ancestral phage stock. The negative values in the Y-axis indicate the number of times the efficiency in the host is lower regarding *Aeromonas*. The second column (C and D) indicates the trajectories of the difference in EOP as previously described, when the phages sequentially propagated on *Yersinia*. In D, no EOP difference is shown for *Pseudomonas* (in red) in the time 10 because we could no longer count plaques of the phage 23 on this host to estimate its EOP. Finally, in the third column (E and F) a rotative host propagation strategy was used, where the phages were alternatively replicated in *Pseudomonas* SR91, *Yersinia* XS2-2 and *Aeromonas* XB1-6. R1, R2 and R3 represent independent lineages from the same ancestral stock (i.e. phage populations initially genetically identical) supported by 5 technical replicates each. The filled circles show the averages and the bars represent the standard deviation for each lineage. All the data plotted in the charts are presented in the Supplementary Table 5.

In parallel to the serial replication in a single strain, we also propagated the Atoyac phages cyclically alternating their host between *Yersinia, Pseudomonas*, and *Aeromonas* to explore whether this condition alters the dynamics in EOP resulting from continuous replication using a single host (Supplementary Figure 7, scheme C). We found that changes in EOP for both phages over the course of the experiment resembled those recorded when the phages are exclusively propagated in the *Pseudomonas* strain, or when Atoyac1 is continuously replicated in *Yersinia*, i.e. a marked reduction in the EOP gap compared to the reference *Aeromonas* isolate (Figure 5, panels E and F). Interestingly, under this propagation scheme, Atoyac 23 did not lose infectivity towards the *Pseudomonas* strain despite being replicated in the *Yersinia* isolate in four of the ten propagation steps, thus implying that host alternation may be key to maintain promiscuity.

In addition to changes in EOP, we observed variation in plaque morphology in some of the evolved lineages. The Atoyac phages produced similar plaques during the course of the experiment when spotted on bacterial lawns of the *Yersinia* and *Aeromonas* strains but generated conspicuous variants when spotted on the *Pseudomonas* isolate. Atoyac 23 rapidly developed larger bull’s-eye plaque variants when sequentially replicated in *Pseudomonas* (Figure 6, panel B). Similar morphology was also recognized during alternate-host propagation but in most cases was not fixed over the course of the passages (Figure 6, panel F). Remarkably, this new Atoyac 23 plaque morphology in *Pseudomonas* resembles that produced on lawns of the *Aeromonas* isolate, the host with the largest EOP at the beginning of the experiment (Supplementary Figure 2), and contrasts with the morphology observed when the phage is serially replicated in *Yersinia*, where it quickly generates nearly-imperceptible plaque variants (Figure 6, panel D). Atoyac 1 also developed slightly larger plaques but only in the *Pseudomonas*-replicated lineages (Figure 6, panel A).

**Fig. 6.**
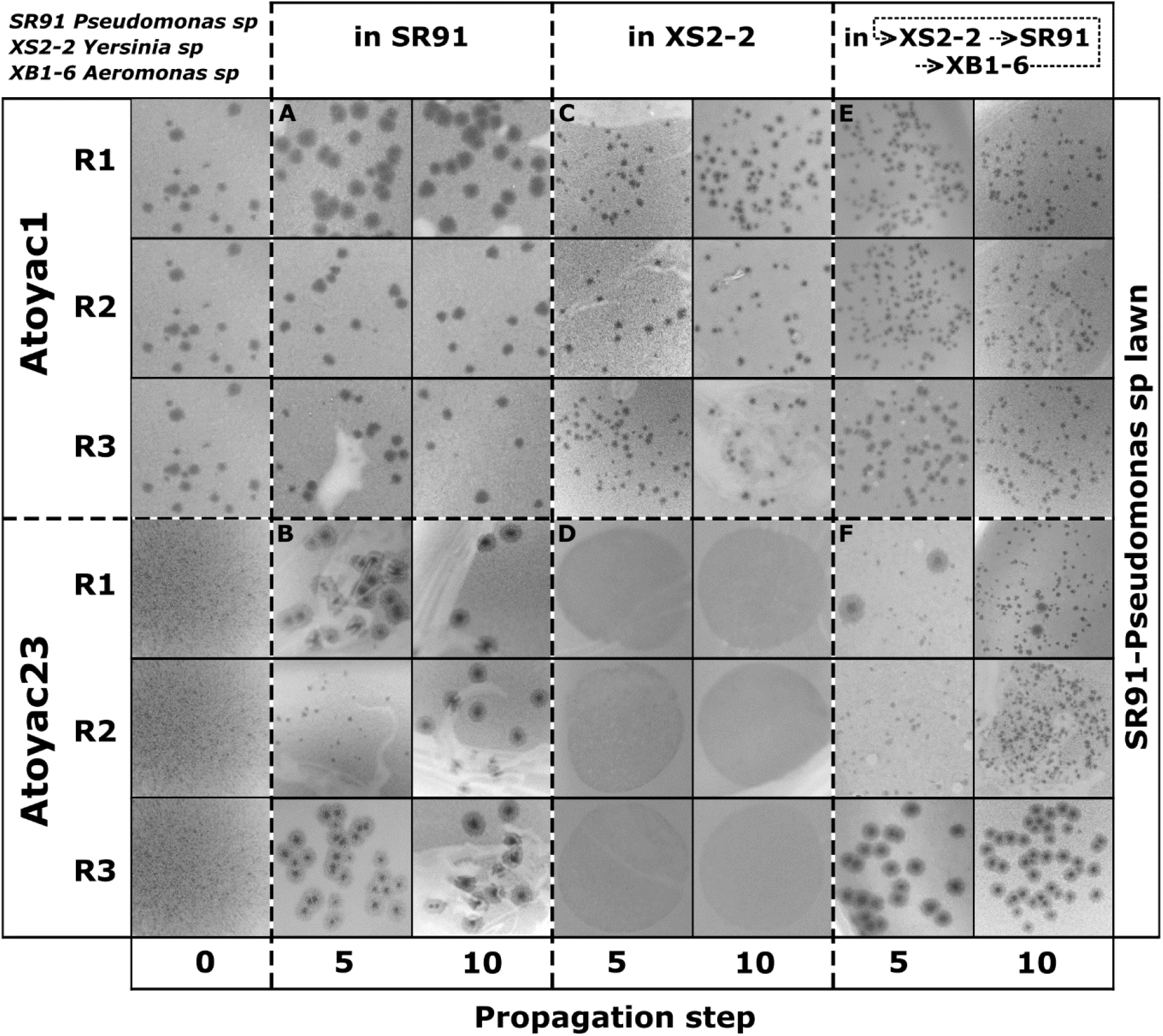
Atoyac phages 1 and 23 plaque morphology comparison on the *Pseudomonas* strain SR91. The figure shows the plaque morphology of the phages Atoyac 1 and 23 in each step of the experimental evolution assay. Three biological replicates (lineages, R1-R3) of the Atoyac phages 1 and 23 were sequentially propagated 5 and 10 times in the *Pseudomonas* strain SR91 (panels A and B), *Yersinia* strain XS2-2 (panels C and D) or alternately in XS2-2, SR91 and *Aeromonas* strain XB1-6 (panels E and F). The resulting plaque morphologies in the SR91 strain were compared regarding the initial one (first column, propagation 0).

## Discussion

Phages are considered the most abundant and diverse biological entities on earth (16). Although numerous efforts exist to isolate and study new types of phages, a big majority of them are targeted/biased towards isolating phages with a narrow host-range (4), thus limiting our knowledge on phages with infection capabilities crossing between different taxonomic levels. This type of viruses are usually referred to as broad-host-range phages, however, we favor the use of “promiscuous” following the term applied to plasmids with mobilization and replication capabilities crossing multiple taxonomic levels (17). Moreover, the use of the term promiscuous allows the distinction between viruses with hosts of different taxonomic origin and those infecting several strains of the same species, commonly referred to as of broad range (5, 14). Promiscuous phages represent powerful models to further our understanding of phage-bacteria interactions and the evolution of host range, however, it is not yet clear how prevalent they can be (5).

Recent studies predicting phage-host relationships using various strategies have inferred interactions beyond the level of species or genus, suggesting that the abundance of promiscuous phages may be higher than currently postulated (4, 5, 7, 8). Previous attempts to isolate promiscuous phages primarily relied on enrichment methods using multiple bacteria of different species and mainly differ in either simultaneous or sequential addition of the potential hosts. These approaches led to the identification of phages infecting strains of two different genera or multiple species within a genus (9, 10). In these reports either ATCC or other well-known type strains (e.g. *P. aeruginosa* PAO1 and *E. coli* K-12) were used as hosts (9, 10), which, although proven effective, limits the isolation to phages infecting strains that may not be representative of those existing in different environments. In contrast, our isolation strategy explored the use of a taxonomically diverse collection of bacterial isolates native to the sampled sites as potential hosts (Supplementary Figure 1), under the rationale that isolating promiscuous phages is more likely when multiple bacterial strains used as indicators come from the same microbial communities they occur in (Supplementary Figure 1).

Our isolation strategy allowed the identification of six phages, named Atoyac, infecting bacteria of the *Aeromonas, Hafnia, Yersinia, Escherichia, Serratia*, and *Pseudomonas* genera, which are part of the Gammaproteobacteria class. Following a similar approach regarding the use of native bacteria as potential hosts, other authors were able to isolate a phage from the Antarctic sea infecting strains of two genera from different proteobacteria classes (18).

Detection of phages infecting different species can be the result of a mix of multiple phages that have not been separated properly, leading to misidentification of a promiscuous host range (2). Host range assessment based on bactericidal effect (e.g. lysis spot) rather than productive infection can also lead to its overestimation (2). Here we addressed these issues through comprehensive evaluation of phage purity (Supplementary Figure 8) and determination of host range based on plaque production only. As previously recommended (2), our phage stock preparation included more than three rounds of plaque purification; additionally, the stocks were purified by cesium chloride (CsCl) density gradient prior final host-range determination (Supplementary Figure 1). Our experimental evolution assays also involved ten sequential passages in either single or multiple hosts of different genera (Supplementary Figure 7), conclusively demonstrating integeneric propagation. In our experiments, we observed consistent evidence of phage stock purity (Supplementary Figure 8), including presence of a well-defined band in the CsCl gradients, similar between-phage DNA restriction pattern and genome assembly size, uniformity in plaque morphology, and homogeneity in virion morphology within and between samples.

Our results show that the six Atoyac phages have a similar host range, consistent with their genomic similarity, and indicate that even the most divergent member of the group (Atoyac phage 15, displaying 86% overall sequence similarity to phage 1) is able to infect six bacterial genera. Differences in the EOP on different hosts correlate with the clustering of the phages based on their genomic sequence similarity. We identified a pair of putative structural genes encoding the tail fiber and a glycosidase superfamily protein, and two adjacent regions, one intergenic featuring putative promoters and one encoding a cluster of small ORFs of unknown function, that could account for the differences observed in the infection phenotypes given their sequence variation within the group. Mutations in virion structural proteins, particularly those that are components of the tail, have been reported as responsible for shifts in the host range of several phages, including in experimental evolution studies (19–21). Variable regions encoding accessory ORFs have also been documented in phage genomes (22), however, these have not been extensively characterized in terms of their functional impact on the phage biology.

Although the Atoyac phages infect 6 bacterial genera, the EOP analysis showed that they produce more lytic plaques on *Aeromonas* isolates than in the other susceptible bacteria, suggesting that this phage group is better adapted to infect the *Aeromonas* genus (Figure 2). The highly similar GC content between the Atoyac phages (59%) and *Aeromonas* (58-62% (23)) genomes, and larger lytic plaques formed in the *Aeromonas* strains further support this notion (Supplementary Figure 2). Since the *Aeromonas* genus is known to be abundant in various aquatic environments like rivers (24), we hypothesize it serves as the primary host of the Atoyac phages, whereas isolates from the other susceptible genera, also present in rivers (25, 26), act as secondary hosts. We speculate that the presence of promiscuous phages in rivers may be related to the dynamics of this type of environment which is commonly influenced by strong seasonality (marked seasons of drought and heavy rain) that impact the composition of microbial communities diversifying it regularly (26, 27). Within this context, a promiscuous host range would be advantageous for the phage as it would allow the use of alternative hosts to spread and remain in the community even if the primary host is not within its reach.

We observed statistically significant differences in the Atoyac phages EOP between *Aeromonas* and other susceptible genera but not between *Aeromonas* sub-clusters (Supplementary Table 2), suggesting that infection of distantly-related bacteria involves a biological cost in terms of lytic plaque production. This notion is consistent with observations on other broad-host-range phages that exhibit substantial variation in plating efficiency when infecting distantly-related hosts (9, 10). Variation in the efficiency of transfer or replication between distant hosts has also been reported for promiscuous plasmids (28), implying that the ability to spread amongst multiple taxonomic backgrounds does not come without a cost and that such trade-off is not exclusive of a certain type of mobile genetic element.

The Atoyac phages were isolated from river water samples obtained in different geographic regions across Mexico, suggesting a wide distribution of the group in similar environmental samples. Surprisingly, our search in metagenomics data did not identify sequencing reads mapping to the genome of the phage Atoyac 1. Whilst this outcome suggests absence of the query genome in the explored datasets, it may also be indicative of the low abundance of Atoyac-like phages in the analysed samples. The searching strategy used in this work involves a dataset sub-sampling step to make the search through the SRA metagenomics data more efficient (15) with the cost of being susceptible to some false negatives, particularly when the sequencing reads of certain taxa are in low abundance. Our study shows that culture-based approaches are still a good alternative to get new insights on the virosphere diversity with the additional advantage of leading to the isolation of specimens that can be further investigated in the lab.

The comparative genome analysis identified the Atoyac phages as a novel viral group. Interestingly, the most similar genomes detected outside the group correspond to distantly-related phages infecting either *Yersinia* or *Aeromonas* species, two of the bacterial groups identified as susceptible to infection by Atoyac phages in this work. Although we hypothesize *Aeromonas* as the primary host of the Atoyac phages, they display higher sequence similarity to the genome of the *Yersinia enterocolitica* phage ISAO8 (~48%) than to that of the *Aeromonas salmonicida* phage phiAS7 (~21%). It is unclear whether the phages ISAO8 and phiAS7 are specialized to infect their reported hosts or have a promiscuous host range as the phages of the Atoyac group. Further characterization of ISAO8 and phiAS7 could provide valuable insights into the evolution of the Atoyac phages’ promiscuous host range in the future. Nevertheless, our study highlights the importance of comprehensively characterizing the host range of a phage by including taxonomically-diverse hosts in the panel of tested strains. In the case presented here, the Atoyac phages could have been reported as *Aeromonas* “specific”, given the large number of host strains identified within this genus, thus veiling the broader infection scope of the group.

Isolating promiscuous phages represents an unique opportunity to explore host-range evolution among phylogenetically distant hosts, hence, we exploited this model to investigate the dynamics of infection of two Atoyac phages on non-optimal hosts (*Pseudomonas* and *Yersinia*) using different propagation schemes. Our results show that successive propagation on either single or alternate hosts has a strong effect on the EOP dynamics and its evolutionary trajectory, albeit the phage used is a critical factor in the outcome. In most cases, the serial propagation promoted EOP optimization in the inefficient hosts whilst maintaining the promiscuous host range, thus suggesting this is a well-conserved feature. The propagation of Atoyac 23 exhibited the most contrasting results in the experiment. Continuous replication in *Pseudomonas* allowed the phage to reach an EOP, and develop a plaque morphology, similar to those observed in *Aeromonas*, the initial most efficient host. In contrast, successive propagation on *Yersinia* did not have a substantial impact on the EOP in this host. Such dissimilar outcomes could be associated with the phage initial EOP in both hosts, i.e. Atoyac 23 marked lower efficiency in *Pseudomonas* regarding *Yersinia*, and hence its margin for optimization. This notion is consistent with a recent experimental evolution study of a generalist Cyanophage which acquired a larger number of evolutionary changes with a positive effect on infection efficiency when continuously propagated on a sub-optimal rather than an optimal host (21). Alternatively, evolutionary changes improving EOP in *Yersinia* might require more propagation steps to emerge and fix in the population.

Cross-adaptation and antagonistic-adaptation were observed in our evolution experiments. Serial propagation of Atoyac 1 in *Pseudomonas* improved its EOP in this host as well as in the *Yersinia* isolate, and vice versa. Phage Atoyac 23, in contrast, displayed a substantial reduction in virulence towards *Pseudomonas* as a consequence of its continuous propagation in the *Yersinia* isolate. Replication in alternate hosts prevented the emergence of this effect, highlighting the importance of propagation in multiple hosts to maintain promiscuity in some phages. We speculate that similar propagation patterns could occur in certain environments featuring high microbial diversity, thus leading to preservation of phages with a promiscuous host range. Our results show that the adaptation outcome from continuous propagation on single or alternate hosts is largely dependent on the phage studied, even between highly-similar phages, making the evolutionary trajectories difficult to predict.

## Methods

### Sample collection and coliform count

A dozen samples were collected in 1L sterile flasks from different rivers and sewage in central, northern and southern Mexico. Estimation of water pollution in the different samples was determined by thermotolerant coliform counts on modified mTEC media, following the protocol by membrane filtration previously described (29). Colonies producing a blue pigment (showing Beta-glucuronidase activity) were counted and the CFU/mL were calculated based on the dilution of the samples, in triplicate. Sampling sites with coliform counts of =>1 x 10^3^ CFU/mL were classified as contaminated (Supplementary table 1).

### Bacteria Isolation

A diverse collection of bacterial isolates was assembled from the collected samples. Samples were plated onto different media: LB, BHI, PIA (Pseudomonas Isolation Agar), PAF (Pseudomonas Agar F), PAP (Pseudomonas Agar P), NAA, mTEC, mFC, TSI, and MC (MacConkey), and then incubated for 24h at 30°C (30). After incubation, some colonies were selected somewhat randomly, but looking for diversity in colony morphology, size, and colour, and purified by 3 consecutive passages on the same medium they were isolated from. Pure isolates were grown overnight in BHI broth at 30°C and stored at −80°C in 30% glycerol. The final collection was composed of around 600 isolates coming from 12 different sampling sites.

### Phage Isolation

Bacterial lawns of the collection of isolates were made by mixing 80-250 μL of overnight cultures with 3.5 mL of top agar (1% peptone, 0.5% NaCl, 0.7% Bacto agar, 10mM MgSO4) and plating them on TΦ (1% peptone, 0.5% NaCl, 1.1% Bacto agar), LB, BHI, mFC, TSI, MC or MC without salts media. Aliquots of the environmental water samples used to isolate bacteria were centrifuged and filtered (0.22 μm Millipore membrane) and then spotted on the lawns in order to identify the bacterial isolates that could be infected by the phages present in the samples. Lawns of the susceptible isolates were then used to spot dilutions of the water samples to visualize isolated plaques that indicate phage infection. In order to aid the visualization of the lytic plaques, rosolic acid (0.01%), phenol red (0.0024%), or bromophenol blue (0.0024%) were used in some of the media.

Cyclic and simultaneous cross infection assays were then used to identify candidate promiscuous phages (i.e. phages that could infect a wide diversity of isolates). Briefly, individual plaques were picked with a sterile toothpick and streaked onto bacterial lawns of the susceptible isolates either simultaneously (all isolates at once) or sequentially (from one isolate to the next one) (Supplementary Figure 1). Phages infecting multiple isolates were considered as candidates to promiscuous phages. These phages were then purified by 4 sequential passages in the strain PIA_XB1_6 as it was overall the optimal host. Additional purification passages were avoided to reduce a possible domestication effect of phages in this host. High titre stocks were generated from the propagation of the last purification pass as described previously (22).

### Taxonomic identification of bacterial isolates to genus level

Bacterial isolates that were susceptible to infection by phages were grown on LB overnight at 30°C. Genomic DNA was extracted from the cultures by following the protocol previously reported by (31). The extracted DNA was used as template for PCR-amplifying the genes gyrB and rpoD.

Oligonucleotides to amplify the gyrB (Forward 5’-GGCGGTAARTTYGAYGATAAC-3’, Reverse 5’-TAGCCTGGTTCTTACGGTT-3’, reported here) were used in those isolates that displayed phenotypic traits similar to the Enterobacteriaceae family (e.g. growth in MC plates). Primers flanking rpoD (Forward 5’-ATYGAAATCGCCAARCG-3’, Reverse 5’-CGGTTGATKTCCTTGA-3’ (32)) were used with the rest of the bacterial isolates. PCR assays were performed using Thermo Scientific Taq DNA polymerase with 0.2 μM each primer and the following cycling conditions: 95°C 3’, 30 × (95°C 30”, 52°C 1’ for gyrB and 56°C 1’ for rpoD, 72 °C 1’), 72 °C 10’. PCR products were purified using MultiScreen^®^ PCRμ96 and sequenced by Sanger sequencing in Macrogen, Inc. The bacterial genera were determined by performing a BLASTn of the obtained sequences against the NCBI database in order to identify the genera of the closest matches. The homologous sequences (using only type strains as reference) were aligned using Clustal omega and used to generate Neighbor-Joining trees (33).

### Promiscuous phage purification

Phage stocks were treated with DNase I and RNase (1 μg/mL) and incubated for 30 min at 37°C. Phage particles were then precipitated with 16% w/v PEG 8000 and 1.4M NaCl overnight at 4°C, and pelleted at 8,000 G for 20 min. The supernatants were discarded and the pellet was resuspended in 1 mL of SM buffer (50 mM TrisHCL-pH 8, 10 mM MgSO4, 100 mM NaCl, and 0.01% Gelatine). In order to remove the PEG from the stocks, an equal volume of chloroform was added to the tubes and centrifuged at 8,000 G for 5 min. The aqueous phase was recovered and purified by ultracentrifugation in a discontinuous CsCl gradient, as previously described (34). Finally, the pure phage stocks were dialyzed to remove the salts, and stored at 4°C.

### Determination of phage host range and efficiency of plating

Cross infection assays (see above) were used to characterize the host range of the promiscuous phages based on their ability to produce isolated plaques (productive infections) in the different bacterial isolates. Phages that were capable of infecting different taxonomic genera were classified as promiscuous and named Atoyac phages which means river in Náhuatl, an indigenuous language of Mexico. To determine the efficiency of plating (EOP) of the Atoyac phages, serial dilutions of the CsCl-purified stocks were spotted onto the susceptible isolates of the bacterial collection. The EOP was calculated as the average of the titre of the phage in a given strain divided by the titre in the isolate that was used to initially propagate the phage (PIA_XB1_6). Each infection assay was performed with 5 replicates.

### Statistical analysis

In order to determine whether the data of efficiency of plating was normally distributed, histograms, quantile-quantile (QQ) plots and Shapiro-Wilk tests were used taking as input the EOP values of each phage on the different bacterial genera. As the data was not normally distributed due to the nature of the samples, the non-parametric Mann-Whitney U test was used and p-values were corrected for multiple comparisons using the Bonferroni correction. Corrected p-values lower than 0.05 were considered as statistically significant (Supplementary Table 2). For the experimental evolution data, one-way ANOVA tests were used to compare the difference of EOP for each condition among the different time points. p-values lower than 0.05 were considered as statistically significant (Supplementary Table 5).

### Virion morphology

The virion morphology was determined by negative contrast stain transmission electron microscopy as previously described by (22). Briefly, aliquots of the pure phage stocks were deposited on a carbon-coated copper grid and stained with 2% uranyl acetate. Grids were examined under a JEM-2000 transmission electron microscope at 80 Kv.

### Phage DNA extraction and genome sequencing

DNA was extracted from 500 μL of CsCl-purified phage stocks following the protocol previously described (34), excluding the step with proteinase K. Values of purity and concentration were measured by spectrometry (Nanodrop 2000). Aliquots of 500 ng of DNA were used to determine the size of the genome by restriction digestion. Finally, the DNA for the different phages was sequenced by Illumina (pair-end) technology in the UUSMB-UNAM.

### Phage genome analysis

The reads were adapter-trimmed and quality-filtered using Trimmomatic v.0.36 (35). The filtered reads were then assembled using Velvet v.1.2.10 (36) and mapped against the de novo assemblies using BWA v.0.7.17 (37). The files with the mapped reads were then used to correct the assemblies using Pillon v.1.22 (38). The genomes were preliminary annotated with Prokka v.1.13 (39) and re-annotated manually based on protein homology searches and conserved domains using InterProScan 5.30-69.0, Psi-BLAST and CD Search (40–42).

The genomes were used for comparative genomics analysis of the Atoyac phages and their homologous, 4vB_YenP_ISAO8 (NC_028850.1) and phiAS7 (JN651747.1) phages. The Coding Sequence (CDS) regions of Atoyac1 phage were used as reference to compare with the other phages using the GView Server (https://server.gview.ca/ (43)), and generate a comparative circular map (BLAST Atlas). Similarly, Atoyac phages were aligned using Mauve v.2.4.0 with the default parameters to create a comparative linear map and closely visualize the main variable regions of the genomes (44). Finally, the phages’ genomes were globally aligned with Clustal omega and a BioNeighbor-Joining tree was constructed based on the alignment (33).

### Promoters prediction

The *In Silico* promoters search for the intergenic region was performed using the phage Atoyac 1 sequence (from nucleotide 21377 to 23783), as reference of the group, with the promoters prediction tools (for both strands): Neural Network Promoter Prediction (NNPP), PromoterHunter and BPROM through of their web servers (http://www.fruitfly.org/seq_tools/promoter.html, ww.phisite.org/main/index.php?nav=tools&nav_sel=hunter, www.softberry.com/berry.phtml?topic=bprom&group=programs&subgroup=gfindb (45–47)).

### Search of Atoyac genomes in metagenomics data

The genome of phage Atoyac 1 was used as a reference to search for homologues in metagenomics data deposited in the NCBI sequence read archive (SRA). Whole shotgun metagenomes available from SRA and previously curated with PARTIE (48) were mapped to the reference genome using bowtie2 (49) with default parameters through the Search SRA Gateway https://www.searchsra.org/ (15, 50, 51). Only alignments of reads > 50 bp were considered. The number of mapped reads and coverage of the alignments was determined from the sorted and indexed BAM files using Samtools v.1.7 (15, 44, 52).

### Experimental evolution of plate efficiency

The experimental evolution strategy consisted of the sequential propagation of the Atoyac 1 and 23 phages (representing different subgroups) by plaque infection assays on a single host, using either the SR91 strain from *Pseudomonas* or the XS2-2 strain from *Yersinia* (which they infect with low efficiency of plating regarding the strain XB1-6 from *Aeromonas*) or on multiple hosts, using alternately the strains XS2-2, SR91 and XB1-6. In both evolutionary schemes, the phages were propagated for ten passages from three isolated plaques (lineages) obtained from the ancestral stock. The efficiencies of plating of phages were calculated at the propagation steps 0, 5, and 10 (using the high-titer stocks generated in these stages) on *Pseudomonas* and *Yersinia*, using the titer in *Aeromonas* as reference, with five technical replicates each. The normalized efficiency regarding *Aeromonas* was obtained by dividing the titer observed on this host by the titer recorded on *Pseudomonas* or *Yersinia* (host X) (Supplementary Figure S7) and plotted as a negative number to represent how many times the infection efficiency in the host X was below the efficiency in *Aeromonas*.

### Sequence Data availability

The Illumina-sequenced genomes and annotations of the Atoyac1, Atoyac10, Atoyac13, Atoyac14, Atoyac15, and Atoyac23 phages were deposited in GenBank under accession numbers MT682386-MT682391.

## Acknowledgments

DC gratefully acknowledges the Programa de Doctorado en Ciencias Biomédicas, UNAM, and the Ph.D. scholarship 586079 from Consejo Nacional de Ciencia y Tecnología (CONACYT, México). PV acknowledges funding from DGAPA-PAPIIT/UNAM IN206318 and CONACYT 23_A1-S-11242. We wish to thank María de Lourdes Rojas-Morales from Microscopía Electrónica, CINVESTAV, for her technical assistance in our electron microscopy studies and Javier Rivera Campos from CCG, UNAM, for technical support with lab experiments.

## Author contributions

DC, AC, WF, and PV conceptualized the study. DC, AC, and WF drafted the manuscript with input from the other authors. DC isolated the phages and carried out the characterization experiments, data analysis and visualization. AC contributed to the genome and metagenome data analysis, the virion purification and characterization by electron microscopy. WF performed the statistical analyses and contributed to the virion purification, phage DNA extraction and characterization by electron microscopy and EOP. GG contributed supervising the work. RE contributed with the metagenomics data analysis and revision of the manuscript. PV was responsible for funding acquisition and contributed to the phylogenomic analysis.

## Materials & Correspondence

Correspondence and material request should be addressed to Pablo Vinuesa or to Daniel Cazares.

## Supplementary information

**Supplementary Figure 1. Diagram of the strategy used for the isolation of promiscuous phages.** 1. Diverse water samples with different levels of pollution were collected. 2. Free phages from these microbial communities were isolated by centrifuging and filtering the water samples. Simultaneously, bacteria from these microbial communities also were isolated to generate a wide and diverse collection of native bacteria. 3. Infectivity of the isolated phages on the native bacteria was evaluated and the susceptible isolates were taxonomically identified. 4. Cyclic and simultaneous cross-infection assays were performed on the susceptible isolates (picking isolated lytic plaques and passing them simultaneously and sequentially through the susceptible isolates). 5. Candidates to promiscuous phages (i.e. those that could infect multiple isolates) were propagated using a single host (4 times) and then purified by ultracentrifugation in CsCl gradients. 6. Finally, the pure phages were subjected to cyclic and simultaneous cross-infection assays to validate their promiscuous host range (for more details, see Methods).

**Supplementary Figure 2. Plaque morphologies of Atoyac phages in representative strains of the different taxonomic groups.** Infection assays for visualization of lytic plaques were performed using CsCl-purified stocks. The different taxonomic groups correspond to those in Figure 1.

**Supplementary Figure 3. Geographical location of the sites where the Atoyac phages were isolated from.** Atoyac phages were isolated from river and sewage samples collected in central (Paraíso-Morelos, Temixco-Morelos, Río de los remedios-CDMX), northern (Río Tijuana-Baja California) and southern Mexico (Paseo de Ovejas-Veracruz).

**Supplementary Figure 4. Electron micrographs of virions of the Atoyac phages 14, 10, 13 and 23.** The phages shown in the figure exhibit the same morphology as the phages 1 and 15, which corresponds to podophages.

**Supplementary Figure 5. Neighbor-joining genomic tree of the Atoyac phages and two homologues.** The tree was constructed from the alignment of the genomes (at the nucleotide level) of Atoyac phages and their two distant homologues. The bootstrap support value (100 replicates) of the branches is shown in each node. The overall sequence identity percentage (i.e. over the entire genome length) of Atoyac phages and their distant homologues (marked with an asterisk) regarding phage Atoyac 1 (reference) is shown in the table, as well as the length and average GC content of each phage.

**Supplementary Figure 6. Genomic comparison of the Atoyac phages and their main variable regions.** A Mauve visualisation of the genomes of the Atoyac phages is shown. Each phage genome is represented by a similarity plot in red, with annotated coding regions shown as white boxes. The height of the plots is proportional to the level of sequence identity in that region, with blank spaces representing sequences that are specific for a particular phage. The two main variable regions among the Atoyac phages are zoomed in in the bottom panel. One of the regions corresponds to two structural ORFs associated with the phage tails (on the left) and the other one corresponds to a non-coding region (enriched in putative promoters-Supplementary Table 3) and to a series of small ORFs of unknown function (on the right). The small blocks of the same color correspond to portions of DNA shared among the phages in this variable region.

**Supplementary Figure 7. Diagram of the strategy used for the experimental evolution assays.**

Atoyac phages 1 and 23 were subjected to two different propagation schemes by plate assays to evolve their efficiency of plating (EOP), one of them using sequential propagation with a single host (A and B) and the other one alternating its propagation between three different hosts (C). CsCl-purified phage stocks (i.e. ancestral stock: propagation step 0) were propagated on *Pseudomonas* (pink lawn) (A) or on the *Yersinia* (grey lawn) (B). After growth, three individual lytic plaques (R1-R3 lineages from the ancestral stock) were randomly picked from the bacterial lawns and propagated independently with a sterile toothpick through 10 passages on *Pseudomonas* (A) or *Yersinia* (B) (propagation scheme with a single host) or cyclilly on *Yersinia, Pseudomonas*, and *Aeromonas* (green lawn) (C) (scheme with multiple hosts). Phage stocks were generated for each of the 3 lineages on the different schemes from propagation steps 5 and 10 and these new stocks of the evolved phages were spotted (as well as the initial stocks corresponding to propagation step 0) on the lawns of *Aeromonas, Pseudomonas*, and *Yersinia* to estimate their EOPs (from 5 technical replicates each) regarding *Aeromonas’* efficiency.

**Supplementary Figure 8. Evidence of the purity of Atoyac phages.** Atoyac phages were isolated following multiple control steps to ensure their purity (for details see Supplementary figure 1 and Methods). A. Stocks purified by ultracentrifugation in CsCl gradient always showed a consistent plaque morphology on the propagative host and other susceptible strains (Supplementary Figure 2). B. A single defined band (containing the pure phages) was observed after the CsCl purification. C. Estimation of the genome size of the Atoyac phages obtained from the digestion of their genomic DNAs with NdeI enzyme (43 450 bp) was very similar to the genome size of the unique assemblies obtained from their genomic sequencing (41.7 kb on average). D. A single virion morphology (corresponding to the podovirus morphotype) was observed through the transmission electron microscope for the pure stocks of Atoyac phages.

**Supplementary Table 1. Sites sampled in this study**

**Supplementary Table 2. Titers and EOPs of Atoyac phages on the susceptible isolates and Mann-Whitney U test of EOP differences among taxonomic groups and subgroups by each phage**

**Supplementary Table 3. Promoters prediction of the intergenic region**

**Supplementary Table 4. Metagenome reads mapping the crAssphage and T4 phage genomes**

**Supplementary Table 5. EOPs from experimental evolution assays and ANOVA test**

